# A Cell Culture Model of BK Polyomavirus Persistence, Genome Recombination, and Reactivation

**DOI:** 10.1101/2021.08.05.455229

**Authors:** Linbo Zhao, Michael J. Imperiale

## Abstract

BK Polyomavirus (BKPyV) is a small non-enveloped DNA virus that establishes a ubiquitous, asymptomatic, and lifelong persistent infection in at least 80% of the world’s population. In some immunosuppressed transplant recipients, BKPyV reactivation causes polyomavirus-associated nephropathy and hemorrhagic cystitis. We report a novel in vitro model of BKPyV persistence and reactivation using a BKPyV natural host cell line. In this system, viral genome loads remain constant for various times post-establishment of persistent infection, during which BKPyV undergoes extensive random genome recombination. Certain recombination events result in viral DNA amplification and protein expression, resulting in production of viruses with enhanced replication ability.

**Importance:** BK polyomavirus (BKPyV) generally establishes a persistent subclinical infection in healthy individuals but can cause severe disease in transplant recipients. While an in vitro model to study acute replication exists, no practical model with which to study BKPyV persistence is currently available. We established a BKPyV persistence model in cell culture. Our model reveals that the virus can persist for varying periods of time before random recombination of the viral genome leads to enhanced replication.

## Introduction

Polyomaviruses are a group of small non-enveloped icosahedral DNA viruses about 45 nm in diameter (1). The first polyomavirus was identified in 1953 as a filterable agent that causes salivary gland carcinomas in mice, and two human polyomaviruses were subsequently identified in 1971: BK polyomavirus (BKPyV) was isolated from a kidney transplant patient with the initials B.K. who was hospitalized for a ureteric obstruction (2) and JC polyomavirus was cultivated from a progressive multifocal leukoencephalopathy patient with the initials J.C. (3). Epidemiology studies have shown that BKPyV infection is ubiquitous among the world population, with more than 80% of the population serologically positive for BKPyV (4). After initial exposure, BKPyV establishes a persistent asymptomatic infection in the urinary tract with periodically shedding of infectious virus particles into the urine of healthy individuals (5). However, in some immunosuppressed patients, BKPyV reactivates and replicates to high levels, causing polyomavirus-associated nephropathy in renal transplant recipients and hemorrhagic cystitis in allogeneic hematopoietic cell transplant recipients (6). The mechanisms underlying BKPyV persistence and reactivation are not understood.

BKPyV has a circular double-stranded DNA genome about 5 kb in size, which is divided into three functional regions: early region, late region, and non-coding control region (NCCR; Figure 1A). The NCCR contains the origin of DNA replication along with cis-acting elements that regulate the bidirectional transcription of both early and late genes from opposite strands of the genome (7). There are two genetic forms of BKPyV, which are distinguished by the structure of the NCCR: archetype virus and rearranged variants. Archetype BKPyV is commonly isolated from healthy individuals, and it is regarded as the persistent and transmissible type of virus (8). However, this virus replicates poorly if at all in cell culture (7, 9). The NCCR of archetype virus is arbitrarily divided into five sequence blocks termed O, P, Q, R, and S, while rearranged variants usually have duplications and/or deletions of these blocks (Figure 1B), and it has been shown that the NCCR is the major determinant of replication ability (7, 10, 11). Rearranged variants are commonly isolated from patients with BKPyV disease and can replicate robustly in vitro (12), including in cultured primary human renal proximal tubule epithelial (RPTE) cells (13). Because of a lack of a suitable animal or cell culture model, less is known about infection with archetype virus and the evolutionary processes that lead from archetype virus to rearranged variants. While it is thought that rearranged NCCRs are generated by recombination, direct evidence for this process remains elusive (14, 15).

**Fig. 1.**
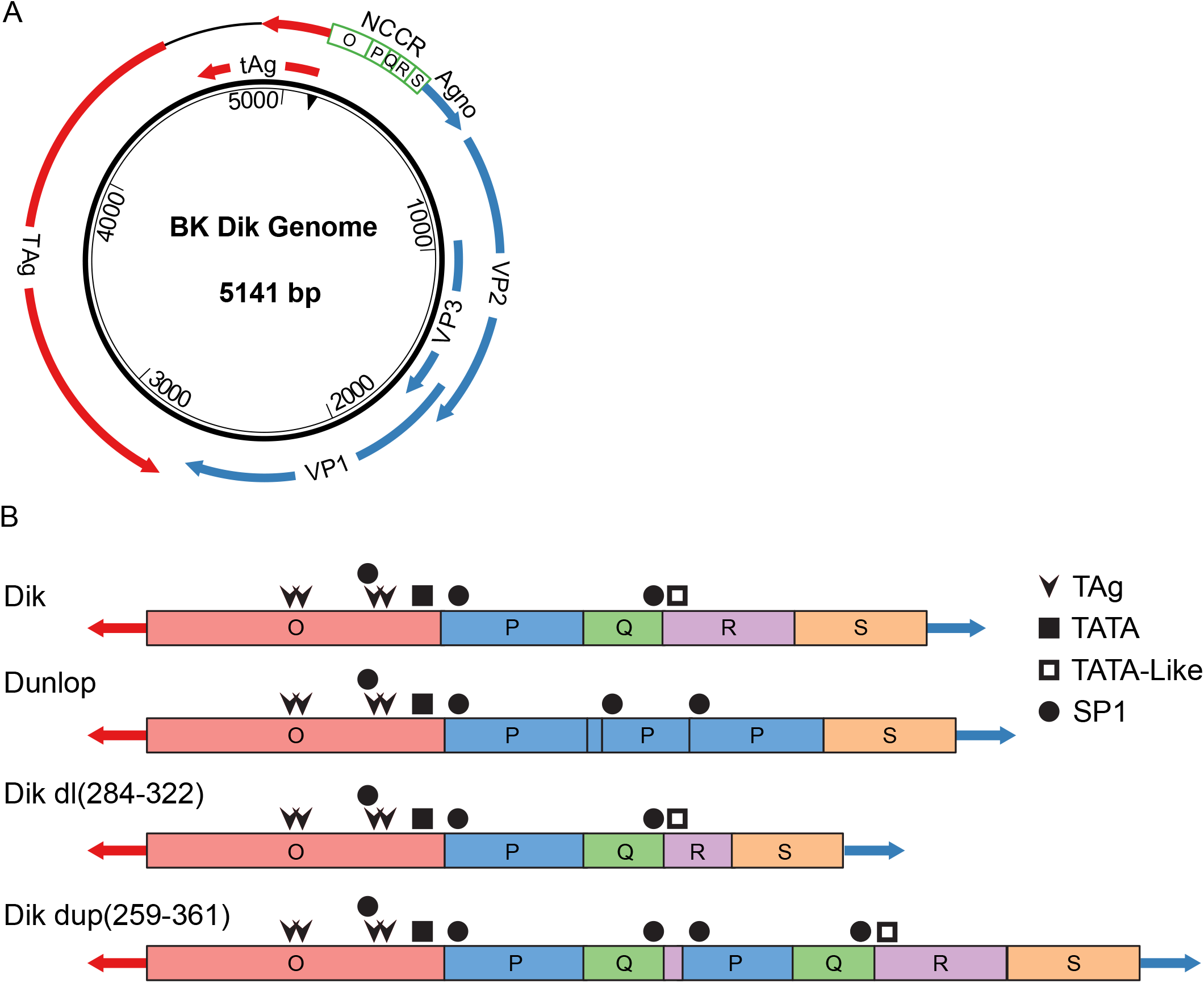
Diagrams of the BKPyV genome and NCCRs. (A) Schematic diagram of the BKPyV genome. Arrows indicate open reading frames. See text for explanation of nucleotide numbering. Color coding of the viral genome is consistent throughout all the figures. (B) The sequence blocks of the archetype (Dik), Dunlop variant, and recombinant NCCRs described in this manuscript are shown. The binding sites for factors discussed in the text (TAg, Sp1), as well as the putative TATA and TATA-like elements, are indicated.

In this study, we established an in vitro model of archetype BKPyV persistence using a human telomerase reverse transcriptase-expressing human RPTE cell line (RPTE-hTERT) (16). This model mimics the immunosuppressed environment in transplant patients in that no adaptive immune system is present. We find that archetype BKPyV can persist in RPTE-hTERT cells for up to 100 days before reactivation and genome amplification occurs. We show that amplification is associated with robust recombination of the viral genome. We carefully mapped the recombination events in BKPyV genomes harvested from different time points during the persistence stage and reactivation. Our next-generation and single-molecule sequencing results indicate that the BKPyV genome continuously evolves with random recombination during persistent infection. Eventually, these recombination events lead to rearrangements in the NCCR that allow increased BKPyV genome replication, gene expression, and production of progeny virus that have enhanced replication ability when used to infect naïve cells. Analysis of the recombination junctions indicates that recombination primarily appears to occur by non-homologous end joining. This model will be useful for future studies of the viral and host factors that mediate persistence and recombination.

## Results

### Archetype BKPyV virus establishes a persistent infection then reactivates in RPTE-hTERT cells

While an in vitro acute BKPyV lytic infection model of RPTE cells with which to study rearranged variants exists (13), primary cells cannot be easily adapted to a persistent infection model for archetype virus due to their limited capability to replicate in culture. As our previous studies showed that BKPyV behaves differently in its natural host cells (RPTE) and non-physiological cell lines such as CV-1 and Vero cells (17, 18), we wished to establish a persistence model by minimally modifying the natural host cells. To overcome the growth limit of primary RPTE cells, we developed a human telomerase reverse transcriptase-expressing RPTE cell line (RPTE-hTERT), which is immortalized but behaves identically to primary cells with respect to lytic infection by rearranged variants (16).

To test if RPTE-hTERT cells can be used as a persistence model for BKPyV, we infected the cells with archetype virus. Cells were passaged and low molecular weight DNA was harvested at various time points post-infection until obvious cytopathic effect (CPE) resulted in an inability to passage the cells further. Viral genome copy numbers were measured by qPCR and normalized to mitochondrial DNA copy number, as a surrogate for cell number (mitochondrial DNA is extracted with low molecular weight viral DNA). This pilot experiment showed that BKPyV persisted at a low genome copy number in RPTE-hTERT cells for 30 days before genome replication and reactivation occurred (Figure 2A). Cells infected with a rearranged variant, however, died in less than a week. To confirm and expand upon this result, we performed seven additional repeats, passaging the cells and harvesting DNA and protein every five days. Our overall results show that BKPyV establishes a persistent infection in RPTE-hTERT cells and maintains a low copy number for 20-100 days before reactivation and genome amplification occurs (Figure 2B). An average kidney cell has approximately 1,000 mitochondria (19); therefore, there are fewer than ten archetype genomes per cell during persistence. Because the cells were passaged at a 1:2 ratio each time yet the BKPyV genome copy level remained stable for up to 100 days, we conclude that BKPyV is replicating at a low level in RPTE-hTERT cells in order to keep up with cell division. After reactivation, the BKPyV genome copy number increased exponentially until CPE and cell death were observed in each experiment.

**Fig. 2.**
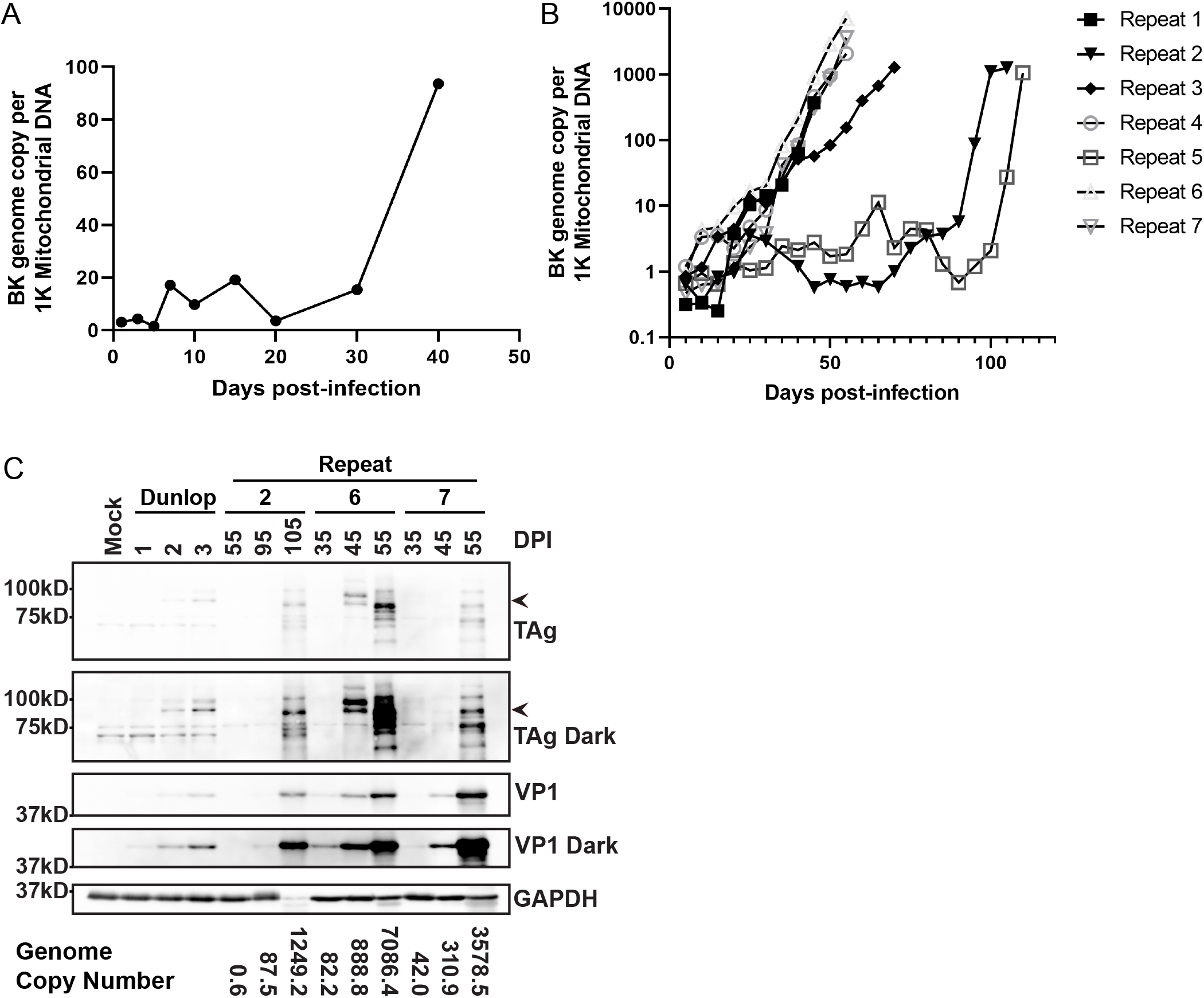
BKPyV genome copy number and viral protein expression during persistence and reactivation. RPTE-hTERT cells were infected at an MOI of 5,000 genomes/cell and samples harvested as described in the text. (A) BKPyV genome copy numbers at the indicated time points post-infection were determined by qPCR and normalized to mitochondrial DNA copy number. (B) BKPyV genome copy numbers for the seven repeats were determined as in A every five days post-infection (C) Protein samples from persistent infection samples and infection with a rearranged variant of BKPyV (Dunlop) were analyzed for viral early (TAg) and late (VP1) protein expression, indicated by the arrows. The DNA copy number for each corresponding sample is indicated at the bottom of the blot. DPI = days post-infection.

To examine viral protein expression during persistence and reactivation, samples collected at representative time points were examined by Western blotting for viral early (large T antigen; TAg) and late (VP1) proteins (Figure 2C). Proteins from infection with a rearranged variant of BKPyV (Dunlop) at 1-3 days post-infection (DPI) were included as a positive control. The results showed that the level of viral protein expression correlates with the genome copy number. TAg and VP1 were barely detectable at 55 days in repeat 2, and at 35 days in repeats 6 and repeat 7. Both TAg and VP1 expression increased as genome copy number increased, indicating progression through the viral life cycle. We also observed variation in TAg protein sizes as the infections progressed, and analysis of the sequences shows examples of deletions that might explain some of the variation (Supplemental Figure S1). These results suggest that BKPyV establishes a persistent infection and eventually reactivates in this in vitro model.

Given the variability in the length of the persistent phase of the infection, we wondered whether we might be able to obtain similar data from an infection of primary cells if robust replication began early. We therefore infected primary RPTE cells in the same manner. In two experiments, we obtained a short enough period of persistence to allow for significant replication before the lifespan limit of the cells was reached (Supplemental Figure S2). These data indicate that the parameters of our system are not an artifact of cell immortalization with hTERT.

### BKPyV genomic recombinants begin to accumulate during persistence

It is generally thought that rearranged variants of BKPyV are derived by recombination of archetype genomes during persistent infection, followed by selection for viruses with enhanced replication ability (14), but this has not been proven experimentally. We therefore sequenced BKPyV genomes over the course of our experiment and examined if recombination was occurring during persistent infection. We prepared DNA libraries directly from low molecular weight DNA with a transposon-based library preparation kit to minimize recombination artifacts that could be generated during the DNA library preparation (20). We also started with a maximum amount of DNA template to minimize the number of PCR cycles required to barcode the library, such that we were able to prepare all libraries with only five PCR cycles. Low molecular DNA from uninfected cells and a plasmid containing the archetype BKPyV genome were included as controls and to validate the recombination identification pipeline. After quality checking and pooling the barcoded libraries, DNA sequences were acquired by reading 250 base pairs from both ends of each DNA fragment. 300-600k reads were successfully obtained from each sample. All reads without at least one stretch of 25 bp that matched the archetype viral genome were discarded. The remaining reads were aligned against an archetype genome template using the Basic Local Alignment Search Tool (BLAST), and analysis was performed as described in the Methods to identify recombination events. Out of more than 470,000 reads from each negative control, we only detected 3 apparent recombination events in our uninfected control and 247 false recombination events (∼0.05% of the total reads) in our plasmid control. These false-positive events are probably due to pooled sequencing result parsing errors and low levels of artifactual recombination introduced by PCR during the library preparation step in the plasmid sample. The overall low false-positive rate suggests that our library preparation protocol and recombination detection assay were robust and not prone to artifacts. In the experimental samples, we detected a very large increase over time in the number of reads containing recombined segments of the viral genome. During the exponential replication stage, 6.7%-14.1% of the total reads contained recombined viral segments.

To facilitate the analysis, we took the Dik genomic sequence and arbitrarily reassigned the number 1 to nucleotide 5034 in the GenBank sequence, which is the first nucleotide in the NCCR (Figure 1A). Thus, the sequence we used starts at the early region junction of the NCCR and ends with the TAg start codon on the opposite strand. The results from the original experiment and three representative experiments from the seven additional repeats are depicted as circular diagrams in which the recombination junctions are linked with lines (Figure 3A, Row 1; Supplemental Figure S3). The intensity of each line reflects the number of reads corresponding to that junction. The data show that there is extensive recombination in the BKPyV genome, and that new recombination events continue to arise as infection progresses.

**Fig. 3.**
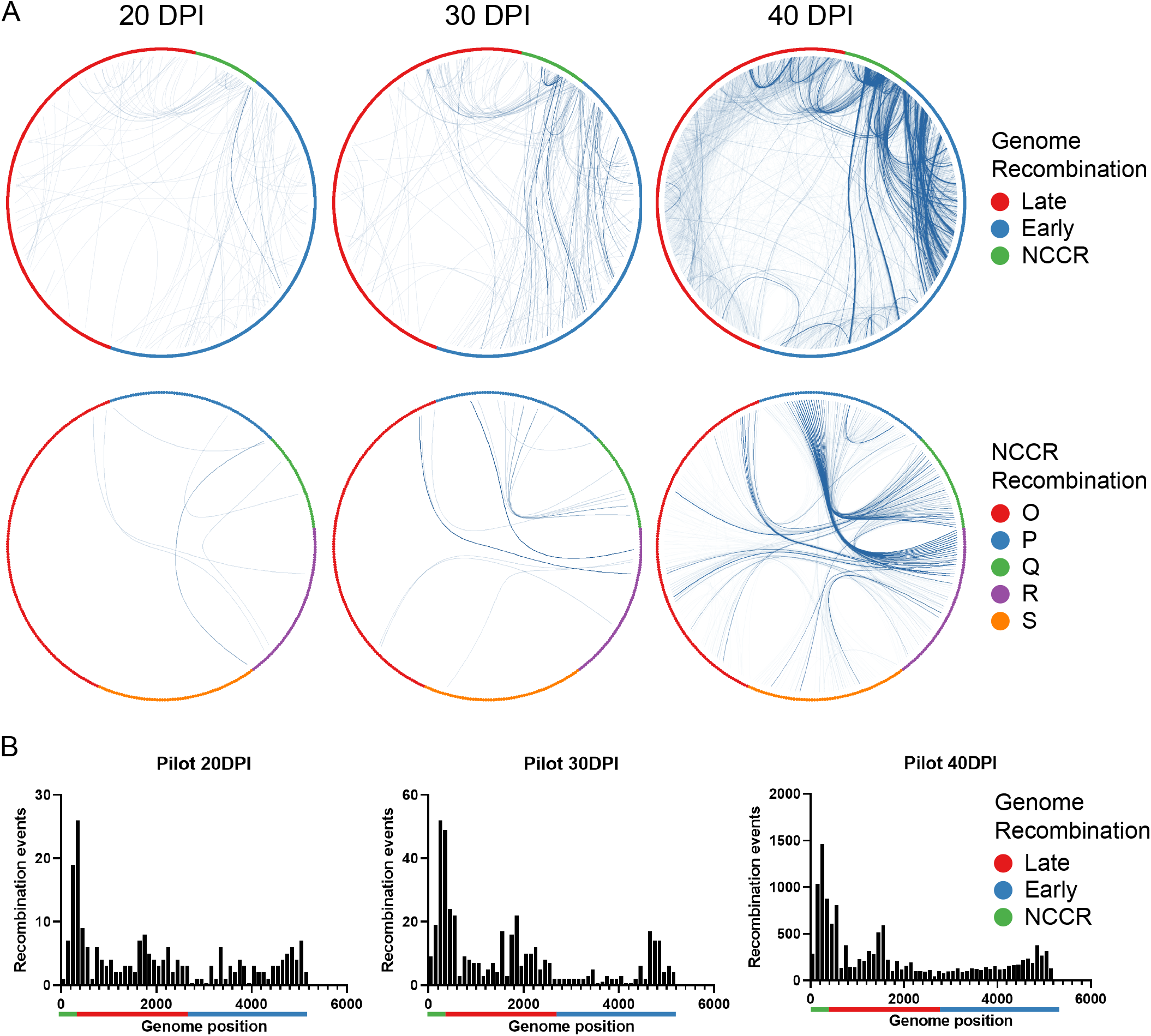
BKPyV genome recombination during persistence and reactivation. (A) BKPyV recombination junctions within the whole genome (Row 1) or NCCR (Row 2) at the indicated time points from the pilot experiment are depicted as circular diagrams in which the junctions are linked with lines. The 40 DPI diagrams were generated with long read sequencing. (B) Distribution of unique recombination events. The distribution of recombination events at representative time points is illustrated as a histogram, with each bar reflecting 100bp. Note the different y axis scales. X-axis legend: Green bar: NCCR. Red bar: late region. Blue bar: early region.

To view recombination events within the NCCR in greater detail, we also graphed them in the circular format (Figure 3A, Row 2; Supplemental Figure S4). These diagrams show that recombination in the P and Q blocks appears to be enriched, suggesting that P and Q block recombination is more likely to benefit viral genome replication, and which is consistent with previous analyses of NCCR function (21). In addition, we graphed the distribution of recombination events as histograms as a more quantitative means of indicating where recombination occurred over time. The histograms show that recombination is fairly evenly distributed in the viral genome at the earlier time points, but at the later time points, genomes with NCCR recombination become more abundant (Figure 3B; Supplemental Figures S5, S6). These results suggest that the BKPyV genome accumulates a significant amount of recombination events and evolves rapidly.

### BKPyV recombination enhances viral replication

To address the relationship between recombination and viral genome replication, we identified an abundant recombinant NCCR sequence and studied its effect on viral replication. To do this, we took an independent DNA sample that had been isolated at 40 days post-infection and amplified the NCCR using PCR. The products were separated on a non-denaturing polyacrylamide gel, and the two most abundant bands were isolated. After sequencing the PCR products, we determined that one band represented the starting archetype NCCR but the other contained a 39 bp deletion in the R and S blocks of the NCCR [Figure 1B, Dik dl(284-322); Figure 4A, red line]. According to our deep sequencing data, this deletion constitutes 3.45% percent of all NCCR recombinant reads at 40 days post-infection. To test the effect of this deletion on replication, we substituted it for the wild-type NCCR of the archetype genome. After growing this virus in 293TT cells, which complement the replication defect of archetype virus (22), RPTE cells were infected with both archetype virus and the archetype virus with the deletion [Dik dl(284-322)]. We harvested low molecular weight DNA and assayed DNA replication by qPCR. The results show a significant increase in replication in the Dik dl(284-322) virus compared to archetype virus (Figure 4B). To examine viral protein expression, RPTE cells were infected with a rearranged variant (Dunlop), archetype virus (Dik), and the Dik dl(284-322) virus. Western blotting shows that the rearranged variant robustly expresses TAg and VP1 proteins, while the archetype virus does not express a detectable level of either protein (Figure 4C). Dik dl(284-322) expresses both proteins at an intermediate level, consistent with the DNA replication data. This result indicates that recombination during persistent infection generates NCCR sequences that allow enhanced BKPyV protein expression and replication.

**Fig. 4.**
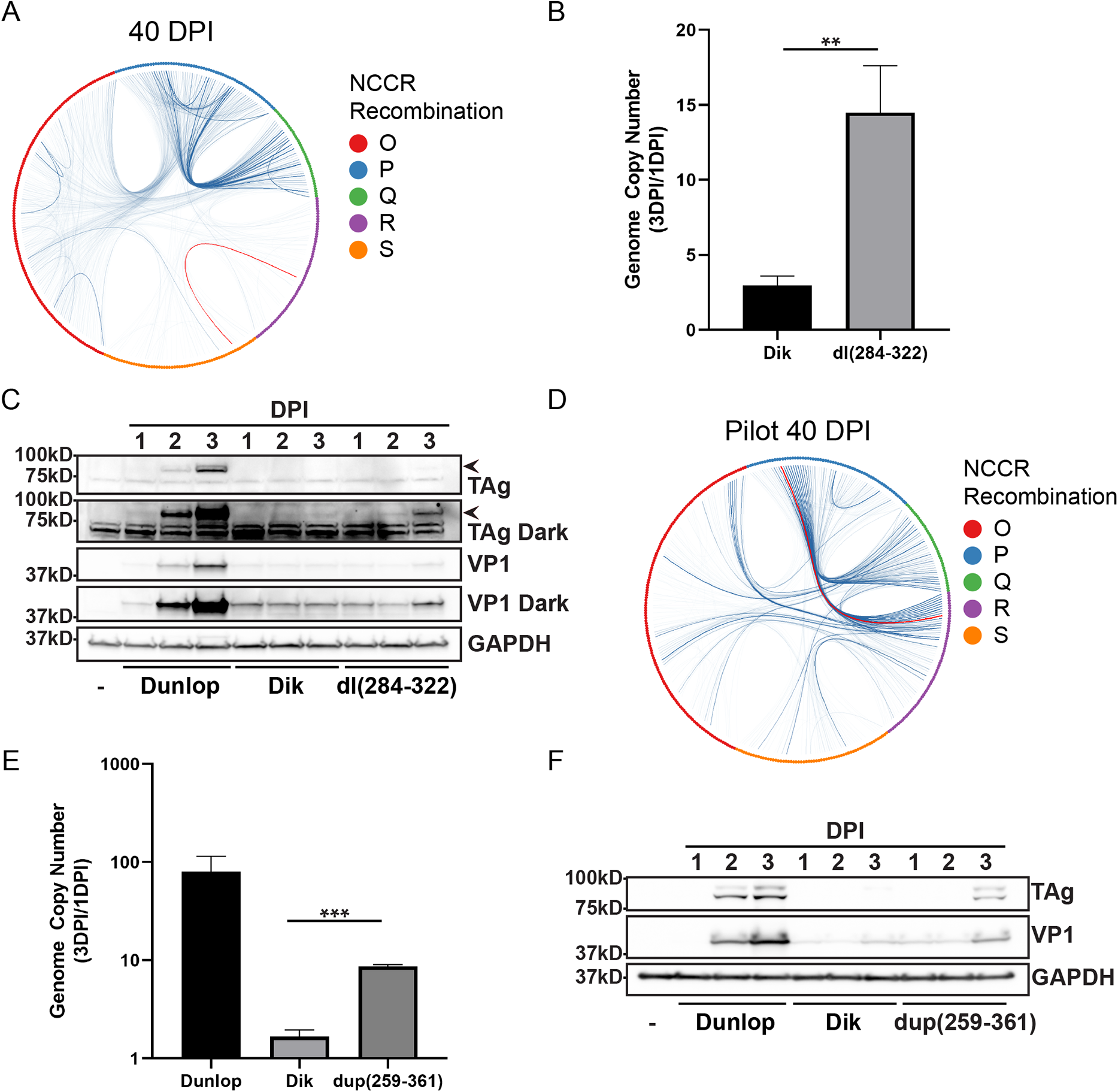
Genome replication and viral protein expression of wild type and recombinant BKPyV. (A) BKPyV recombination junctions within the NCCR at 40 DPI are depicted as circular diagrams in which the junctions are linked with lines. The intensity of each line reflects the number of reads corresponding to that junction. The red event line represents the deletion in Dik dl(284-322). (B) Virus genome replication. Cells were infected with BKPyV wild type (Dik) and the recombinant Dik dl(284-322), and genome copy numbers were measured at 1 and 3 days post-infection (DPI) and normalized to mitochondrial DNA (Mean ± SD; ** P ≤ 0.01). (C) Viral protein expression. Protein samples from infection with BKPyV Dik, Dik dl(284-322), or Dunlop infection were assayed for TAg (arrow) and VP1 expression. (D) NCCR circular diagram at 40 DPI. The red event line represents dup(259-361). (E) Virus genome replication. Cells were infected with BKPyV Dunlop, Dik, or Dik dup(259-361), and genome copy numbers increase were measured at 1 and 3 DPI and normalized to mitochondrial DNA (Mean ± SD; *** P ≤ 0.001). (F) Viral protein expression. Protein samples from the three viruses were assayed as in (C).

### Long-read single molecule sequencing reveals complex and lethal recombination events

Since full-length complete BKPyV genomes could not be assembled from the short-read sequencing results, we turned to long-read sequencing to examine complete recombinant genomes. Low molecular weight viral DNA from infected RPTE-hTERT cells during the exponential replication phase in the pilot experiment (Figure 3A, right column) was amplified with rolling circle amplification (RCA). The RCA products were digested with BamHI to release linear BKPyV genomes and the sequences of these genomes were read using the Nanopore MinION system. We did not detect any significant evidence of genomes that were not monomeric. The reads were aligned against an archetype genome template using BLAST. The results confirmed our observation that BKPyV accumulates extensive recombination events when replication begins. To characterize an individual genome, we cloned a dominant RCA product into pGEM7. This genome has a duplication of the P and Q blocks, and a partial duplication of the R block [Dik dup(259-361)] (Figure 1B; Figure 4D, red line), but no other mutations within the rest of the genome. Our sequencing results showed that the dup(259-361) first appeared at 15 days post-infection. We propagated the Dik dup(259-361) virus in 293TT cells and examined its replication ability as we did for the Dik dl(284-322) variant above. The DNA replication result showed that there is a 9-fold increase in replication in the Dik dup(259-361) virus compared to the archetype virus (Figure 4E). Western blotting showed that Dik dup(259-361) also expresses TAg and VP1 proteins at an intermediate level between the archetype virus and rearranged Dunlop variant (Figure 4F). These results confirmed that recombination during persistent infection can lead to selection for viral genomes with enhanced replication ability as compared to the archetype virus.

Our single molecule results also demonstrated that multiple recombination events can be identified in a single BKPyV genome, with some of these events deleting and/or duplicating parts or all of viral protein-coding regions. Consistent with this finding, we isolated many recombinant genomes that were not capable of producing progeny in 293TT cells: we could only propagate these genomes as mixed populations (data not shown). In addition, we found a combination of the duplication in Dik dup(259-361) and additional recombination events in single viral genomes.

### Analysis of recombination junctions

We next examined the sequences at the recombination junctions to determine if they might provide an indication of the recombination mechanism. It is extremely difficult to imagine that classical homologous recombination could lead to the types of duplications and deletions that are seen in NCCRs isolated from patients, as well as those that we detected. We therefore assumed that non-homologous end joining (NHEJ) was being used to repair double-strand breaks that are known to occur during polyomavirus DNA replication (23). There are two types of NHEJ, classical (cNHEJ) or microhomology-mediated (MMEJ), both of which are error-prone and can generate deletions and duplications (24). Instead of directly ligating DNA breaks together during cNHEJ, MMEJ takes advantage of a microhomology of less than 25 bp in size between the two parental DNA strands (25). Examples of recombination joints, including those of Dik dl(284-322) and Dik dup(259-361), are shown in Figure 5A. We also graphed the distribution of lengths of homologous sequences at the joints to attempt to distinguish between cNHEJ and MMEJ (Figure 5B). It is noteworthy that such homologous sequences could form from either cNHEJ or MMEJ. To attempt to distinguish the role of cNHEJ and MMEJ, we simulated the distribution of homology lengths that would be generated by ligation of random ends using cNHEJ by computationally fragmenting the BKPyV genome and joining random fragments (Supplemental Figure S7). The range of homology lengths from fifty simulated random annealing experiments are illustrated as red whisker boxes in Figure 5B, and our experimental data match the simulated results. However, MMEJ would yield a similar distribution unless one assumes that higher stretches of homology would be favored during the MMEJ process, which would have resulted in a non-linear decay curve. Finally, we examined the distribution of homology lengths at the junctions over time. The results indicate that the relative frequency of lengths remains constant over the course of the infection (Figure 5C).

**Fig. 5.**
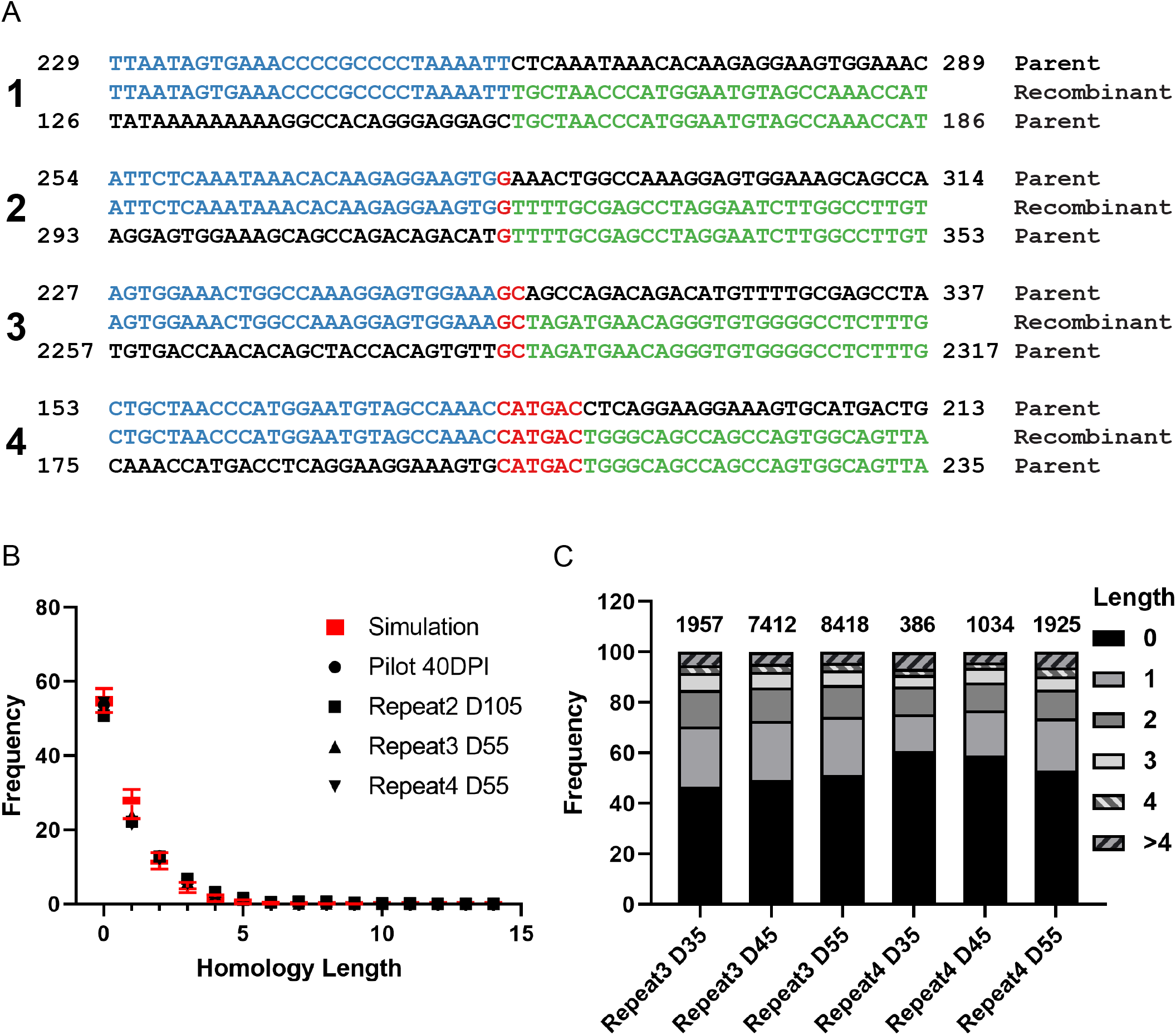
End joining analysis. (A) Representative recombination joints. Blue and green bases indicate the parental DNA fragments, and red bases indicate homology at the joints. Example #1 is Dik dup(259-361) and #2 is Dik dl(284-322). (B) Frequency of homology lengths at recombination joints. The lengths of homology at each unique recombination site were analyzed, and the frequency of each length was plotted (black symbols). The red box plots indicate the simulation as described in the text. (C). Distribution of homology lengths over time. The frequency of lengths at different time points from two representative repeats is shown. The numbers above the bars indicate the number of recombination events analyzed for each bar.

## Discussion

BKPyV establishes a lifelong persistent infection in greater than 80% of the human population after initial exposure during early childhood (4); however, no practical model with which to study BKPyV persistence is currently available. A squirrel monkey model of BKPyV was reported in 2005 (26), but no subsequent studies adopted this model, likely because of the expense of research using non-human primates. Most in vitro models developed for BKPyV have focused on acute lytic infection. Our lab previously reported that RPTE cells could be applied to these studies for rearranged variants of BKPyV (13), and various labs now employ this model to study aspects of viral biology including the viral life cycle, the innate immune response, and the DNA damage response (27-35). However, RPTE cells cannot be used routinely for persistence studies because these primary cells start to grow slowly at passage seven and stop replicating by passage ten, which limits the capability to use this primary cell line to study persistence for prolonged periods.

To establish a robust in vitro model of BKPyV persistence, we therefore extended the life span of RPTE cells by introducing hTERT to establish the RPTE-hTERT cell line (16). Our results show that archetype BKPyV genome copy number remains stable in these cells for up to 100 days before exponential genome amplification starts. Thus, archetype BKPyV successfully established a persistent infection in these cells. Because we harvested and diluted the cells every five days, the BKPyV genome must replicate at a steady low level to maintain a stable copy number. On the other hand, infection of RPTE-hTERT cells with an equal inoculum of rearranged virus results in death by day 5 (data not shown). Western blotting showed that viral TAg expression is not detectable during the persistent stage (Figure 2C), but we cannot rule out that low-level TAg is required for genome maintenance. It is possible that the outcome in infection differs from cell to cell, as had been recently reported (36). However, we think that it is difficult to explain the varying length of persistence if this were the case. Persistence does not seem to be an artefact of hTERT expression since we were able to detect it in primary cells.

As the infection progressed in RPTE-hTERT cells, we detected an eventual exponential increase in BKPyV genome copy number that led to visible CPE and cell death. The time at which this occurred varied from repeat to repeat, which is consistent with a model in which recombination occurs randomly and leads to a replication-competent rearrangement(s) at different times in each repeat. Our deep sequencing and single molecule sequencing results confirmed that the population of recombinants differs in each repeat. We also detected viral early and late protein expression when the genome amplified but not during the persistent phase. In addition, Western blotting indicated variations in the size of the TAg protein after genome amplification began, which suggests recombination occurred in the TAg gene. Indeed, this recombination within the early region was detectable in both the short read and whole-genome sequences, and may lead to changes in the length of the open reading frame or cryptic splicing of mRNA transcripts.

The deep sequencing results showing a high level of genome recombination occurring during persistent infection is consistent with clinical observations that variation exists in polyomaviruses isolated from patients with polyomavirus-related disease (11, 37-39). We detected new recombination events as persistent infection progressed, with most of the recombination in the NCCR (Supplemental Figure S5, S6). This indicates that recombination in the NCCR provides an advantage for viral genome replication and outgrowth, consistent with previous observations that the NCCR is the primary determinant of BKPyV replication (7, 11). We also observed genome recombination outside the NCCR, including some viral-host genome recombination. The recombination outside of the NCCR seems to be less abundant, but this is likely because there is no selection for these events as compared to events within the NCCR. Nonetheless, recombination was random throughout the genome and we did not detect a specific recombination site that is statistically different from the others in the terms of recombination frequency.

At this time, we do not know the mechanism by which the deletion and duplication in dl(284-322) and dup(259-361), respectively, enhance replication. While there are many putative transcription factor binding sites in the NCCR, only Sp1 has been found to actually bind and to play a significant role in the ability of BKPyV to replicate (12, 21). It will be interesting to determine if the recombination events we detected affect transcription, DNA replication, or both.

It is known that BKPyV infection induces a host cell DNA damage response (28, 40). In addition, interfering with the DNA damage response proteins dramatically impairs the quality and quantity of replicated viral genomes, suggesting that BKPyV replication is damage-prone and continuously requires the DNA damage response pathway to resolve replication stress (28). Analysis of replicating viral DNA during infection with the simian polyomavirus SV40 has confirmed that the viral genome is subjected to single- and double-strand breaks. Under ataxia telangiectasia-mutated (ATM) inhibitor treatment, SV40 accumulates strand invasion and rolling circle replication intermediates, while with ATM- and Rad3-related kinase (ATR) knockdown, an increase in broken replication forks can be detected (23). To repair replication stress-associated DNA damage, cells have evolved mechanisms using both homologous recombination and NHEJ. A recent paper described a theoretical model for deletion and duplication in the NCCR (15); however, it cannot fully explain the extensive recombination we detect in the early and late regions of the BKPyV genome. In addition, the NCCR duplications and deletions that have been observed in patient isolates are difficult to explain if homologous recombination were the repair mechanism, and our results indicate that NHEJ, which is more likely to result in small deletions and duplications, is indeed the predominant mechanism.

It is thought that in patients, once rearrangement of the NCCR takes place, the rearranged virus will dominate the population and replace the archetype variant during viral replication. To confirm that rearrangement in the NCCR leads to activation of replication, we isolated a dominant NCCR by PCR amplification of DNA samples harvested after reactivation and used it to replace the archetype NCCR, as well as isolating a virus containing a rearranged NCCR by analysis of complete genomes using single molecule DNA sequencing. Our results showed that a small deletion in the R and S blocks, or a duplication of P, Q, and part of R, leads to a robust increase in viral genome replication, which supports the model that recombination in the NCCR is sufficient to cause genome replication and viral protein expression. This is consistent with our previous finding that the NCCR is the major determinant of the differential replication ability of archetype and rearranged viruses (7). It is also consistent with the observation that many types of NCCR deletions and/or duplications can all lead to enhanced replication (9, 11, 12). It will be interesting to determine how these rearrangements affect transcription of viral genes and the efficiency of DNA replication initiation at the origin.

We also isolated recombined genomes that have sizeable deletions in virus protein-coding sequences. Subsequent experiments showed that these viruses cannot replicate independently, as would be expected: they could be passaged as mixed populations of genomes but did not grow out upon limiting dilution of the viral lysates. However, the fact that these sequences are amplified in the cell suggests that some recombinant genomes may function as helper viruses, with these defective genomes behaving like helper-dependent viruses. This is consistent with clinical findings in which recombinant BKPyV genomes that carry protein-coding sequence deletions can be isolated from urine and kidney allograft biopsies from kidney transplant recipients. These isolated viruses were naturally defective in producing infectious progeny; however, complementation in trans with the missing viral protein could rescue this defect (37).

In this study, we developed a powerful in vitro persistence and reactivation model for BKPyV. We also developed a bioinformatics pipeline for analyzing circular viral DNA recombination. Our model demonstrates that the BKPyV can evolve fairly rapidly, and some of the rearrangements in the NCCR eventually trigger enhanced viral protein expression and DNA replication, and lead to obvious CPE and production of viable viral progeny. Our model mimics an accelerated recombination process in the kidneys of renal transplant patients without an immune system and suggests that BKPyV recombination and enhanced replication are a coordinated process that leads to disease. Our study is the first to demonstrate that recombination can occur throughout the entire viral genome: we propose that most of these recombination events are negatively selected due to their lethality and therefore were not detectable in previous studies examining patient isolates. Our results indicate that BKPyV, even though it is a DNA virus with a low DNA polymerase error rate, can evolve rapidly within an individual host due to recombination. Our model can be used to further study the evolution and reactivation of BKPyV in the kidney, including identification of viral and host factors that contribute to these processes.

## Materials and Methods

### Cell culture

RPTE and RPTE-hTERT cells were maintained in REGM BulletKit media (REGM/REBM, Lonza, CC-3190) at 37°C with 5% CO2 in a humidified incubator (16). Cells were passaged by detaching with 0.25% trypsin-EDTA and split at 1:4 for maintenance.

### BKPyV purification and infection

Plasmids containing the viral genome were digested with BamHI to excise the genome from the vector, and then re-circularized with T4 ligase. The re-circularized genome was transfected into 293TT cells with PEI max transfection reagent (Polysciences, 24765-1). Progeny viral particles were purified on a linear cesium chloride gradient and titered as described previously (27, 41). For infection, cells were pre-chilled for 15 min at 4°C. 1 mL of the virus inoculum was added to one well of a 6 well plate to give an MOI of 5,000 genomes/cell, and incubated at 4°C for 1 hour with gentle shaking every 15 minutes to distribute the inoculum over the entire well. The plate was transferred to 37°C after the 1-hour incubation. As we cannot titer archetype virus using a standard infectious unit (IU) assay due to its inability to replicate, we use genome numbers to normalize different viruses. For reference, 5,000 genomes of a rearranged variant (Dunlop) equals approximately 5 IU.

### Low molecular weight DNA isolation

Low molecular weight DNA was harvested with a modified Hirt extraction protocol developed by the Buck laboratory (42). Briefly, cells were suspended in 250uL Buffer I (50mM Tris, pH 7.5; 10mM EDTA; 50µg/ml RNase A; 20U/ml RNase T1 cocktail) and then lysed by adding 250uL Buffer II (1.2% SDS) and incubating for 5 minutes at room temperature. Cellular DNA was precipitated by adding 350uL Buffer III (3M CsCl; 1M Potassium acetate; 0.67M Acetic acid), incubating at room temperature for 10 minutes, and centrifuging at 16,000xg for 10 min. The supernatant was transferred to mini spin DNA purification columns (Epoch Life Science). 500 uL PB and 750uL PE buffer (Qiagen) were added sequentially to the column and centrifuged to wash the DNA. Low molecular DNA was eluted with 50 uL EB buffer (Qiagen).

### Quantitative PCR

DNA samples were first diluted with DNA grade water at 1:200. Primer pairs amplifying large tumor antigen (TAg) and mitochondrial 16s rRNA gene segments were used as previously described (43). Shapiro-Wilk test, F-test, and Student’s t-test were performed with GraphPad Prism.

### Western blotting

Protein samples were harvested with E1A buffer, electrophoresed, transferred, and probed as previously reported (16).

### Short-read sequencing

The concentration of low molecular weight DNA was measured with a PicoGreen dsDNA Assay Kit (Invitrogen). Sequencing libraries were prepared and pooled according to the manual of the transposon-based Illumina DNA Prep kit (Illumina). 251 bp paired sequences were read using the Illumina MiSeq system with MiSeq Reagent Kits v2. This sequencing was performed at the University of Michigan Sequencing Core.

### Long-read sequencing

Circular low molecular weight DNA was rolling circle amplified with the TempliPhi Amplification Kit (Cytiva). Amplified DNA was digested with BamHI to release linear single genome-length molecules. The digested DNA was separated on a 1% agarose gel and the ∼5 Kb genomic DNA band was harvested with a QIAquick Gel Extraction Kit (Qiagen). The DNA library was prepared according to the manual of the Ligation Sequencing Kit (Nanopore). Long-read DNA sequences were read with the Nanopore MinION system in our laboratory.

### Bioinformatic Recombination Analysis

Step 1. Discard viral DNA reads that do not contain a recombination event. Deep sequencing results (*.fastq.gz) were parsed, and each sequencing read (251 bp in length) that perfectly matches the BKPyV template was omitted from further analysis. Step 2. Discard non-viral DNA reads. Remaining reads were broken into ten 25-mers. If none of the ten 25-mers matched to BKPyV, the original read was omitted from further analysis. All reads that were not omitted were collected to a file in FASTA format for subsequent BLAST. Step 3. BLAST. All sequences collected in the previous step were blasted against the archetype genome sequence with local BLAST+. Step 4. Format BLAST results. Indexes of “start of alignment in query, end of alignment in query, start of alignment in subject, and end of alignment in subject” were parsed from the BLAST results. When a read comprised more than one stretch of DNA originating from different locations of the BKPyV Dik genome, it was considered a recombination event. In addition, the “aligned segment of the query sequence” and “aligned segment of subject the sequence” were parsed from BLAST results, and recombination joints were determined by assembling these two aligned segments as illustrated in Figure 5A. Step 5. Generate recombination diagrams. An R script was used to draw the circular diagrams. Recombination histograms were drawn with Prism. Code used in this report can be found at: github.com/LBZhao/BK_Polyomavirus_recombination_analysis

### Homology length analysis and simulation

After formatting the BLAST result (Step 4, Bioinformatic Recombination Analysis), the homology lengths at the recombination joints were analyzed based on sequence overlap. To simulate homology formation by NHEJ, the BKPyV sequence was first computationally broken into a set of 26-mers, each differing at their 5’ end by one base pair. Random unique 26-mers were religated and the homology length at the center was recorded. If no homology was present at the center, the homology length was recorded as 0. Simulations were performed 50 times with 1000 random annealing events each. Minimum to maximum values are shown with whisker boxes.

### Construction of viral genome with NCCR deletion

An NCCR containing a deletion of nucleotides 284-322 was synthesized with SpeI and SacII sites at the ends and used to replace the archetype NCCR in pGEM-Dik3site (7). The entire genome was confirmed by DNA sequencing. Virus was propagated as previously described and its sequence verified again before use in experiments (22).

### Recombinant viral genome isolation

Gel purified DNA from rolling circle amplification was cloned into pGEM®-7Zf(+)at the BamHI site using T4 ligase. Ligation products were transformed into XL10 cells (Agilent). Colonies containing recombinant genomes were isolated, and DNA was prepared and sequenced. Virus was propagated as previously described and its sequence verified again before use in experiments (22).

## Acknowledgments

We thank Jim Pipas, Adam Lauring, and the members of our laboratory for their critical comments about the manuscript, and Andrew Valesano for advice on single molecule sequencing.

## Author contributions

L. Z.: Conceptualization, Methodology, Software, Investigation, Data Curation, Formal analysis, Visualization, Writing M. J. I: Conceptualization, Methodology, Formal analysis, Supervision, Writing, Funding acquisition.;

## Data and materials availability

Code used in this report can be found at github.com/LBZhao/BK_Polyomavirus_recombination_analysis

## Supplemental figure legends

**Fig S1.**
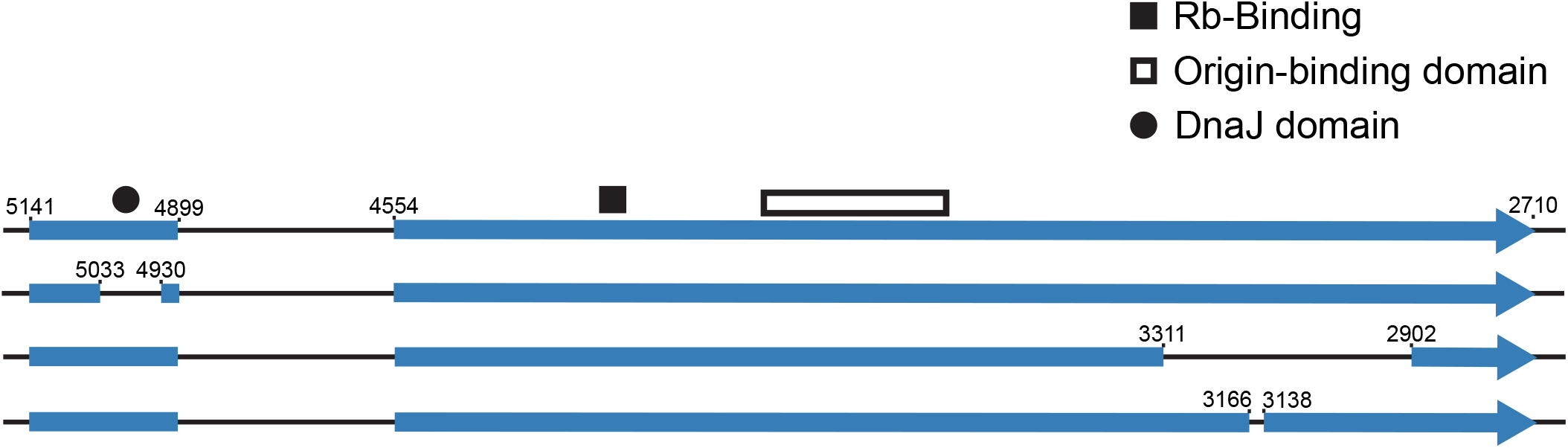
Examples of recombination within the TAg ORF. Representative examples of deletions that might explain the size variation in Figure 2C are shown, along with locations of functional domains (J domain, LXCXE Rb binding domain, origin binding domain) for reference.

**Fig. S2.**
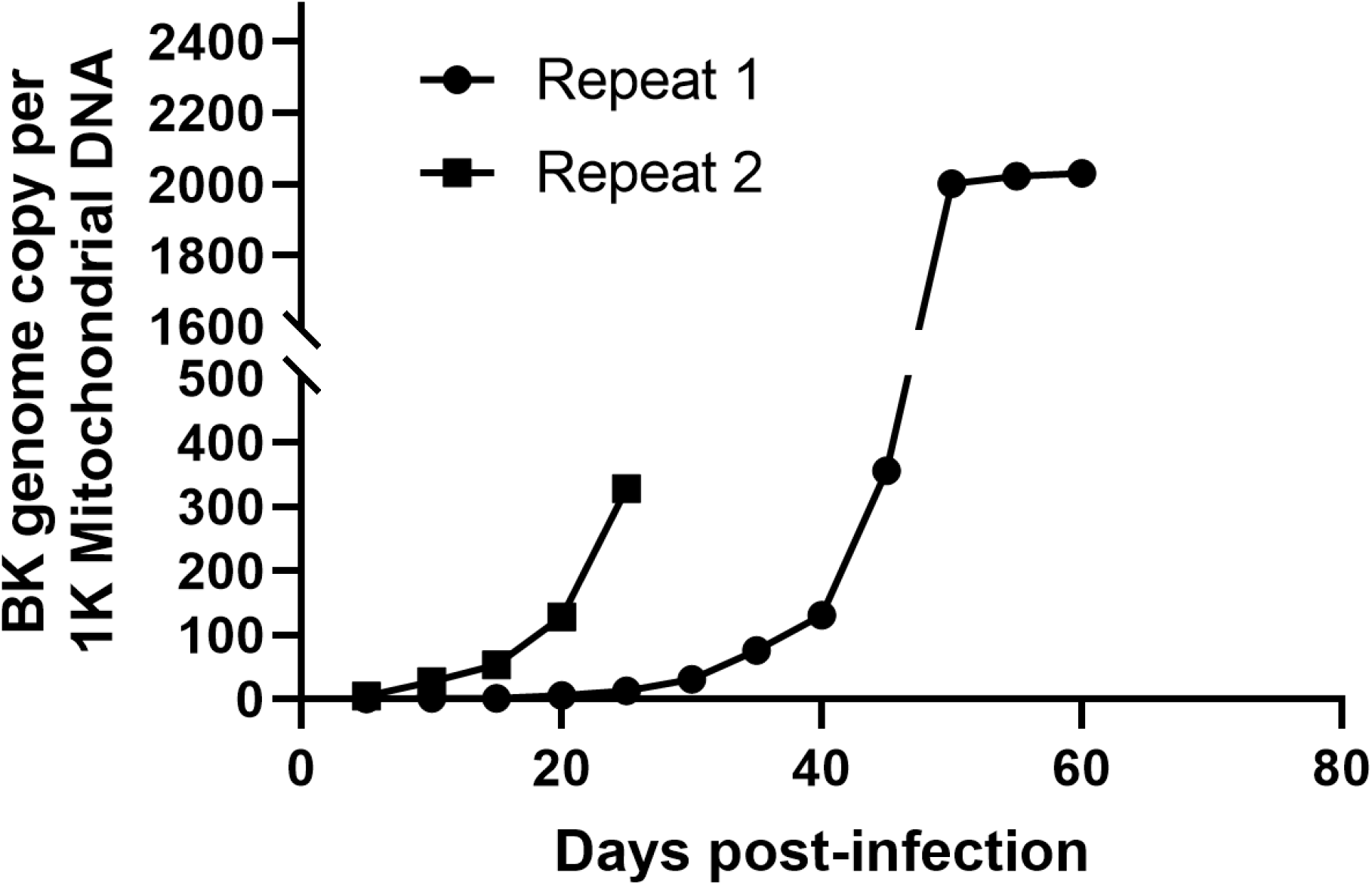
BKPyV genome copy number during persistence and reactivation in RPTE cells. BKPyV genome copy numbers at the indicated time points post-infection were determined by qPCR and normalized to mitochondrial DNA copy number as in Figure 2.

**Fig. S3.**
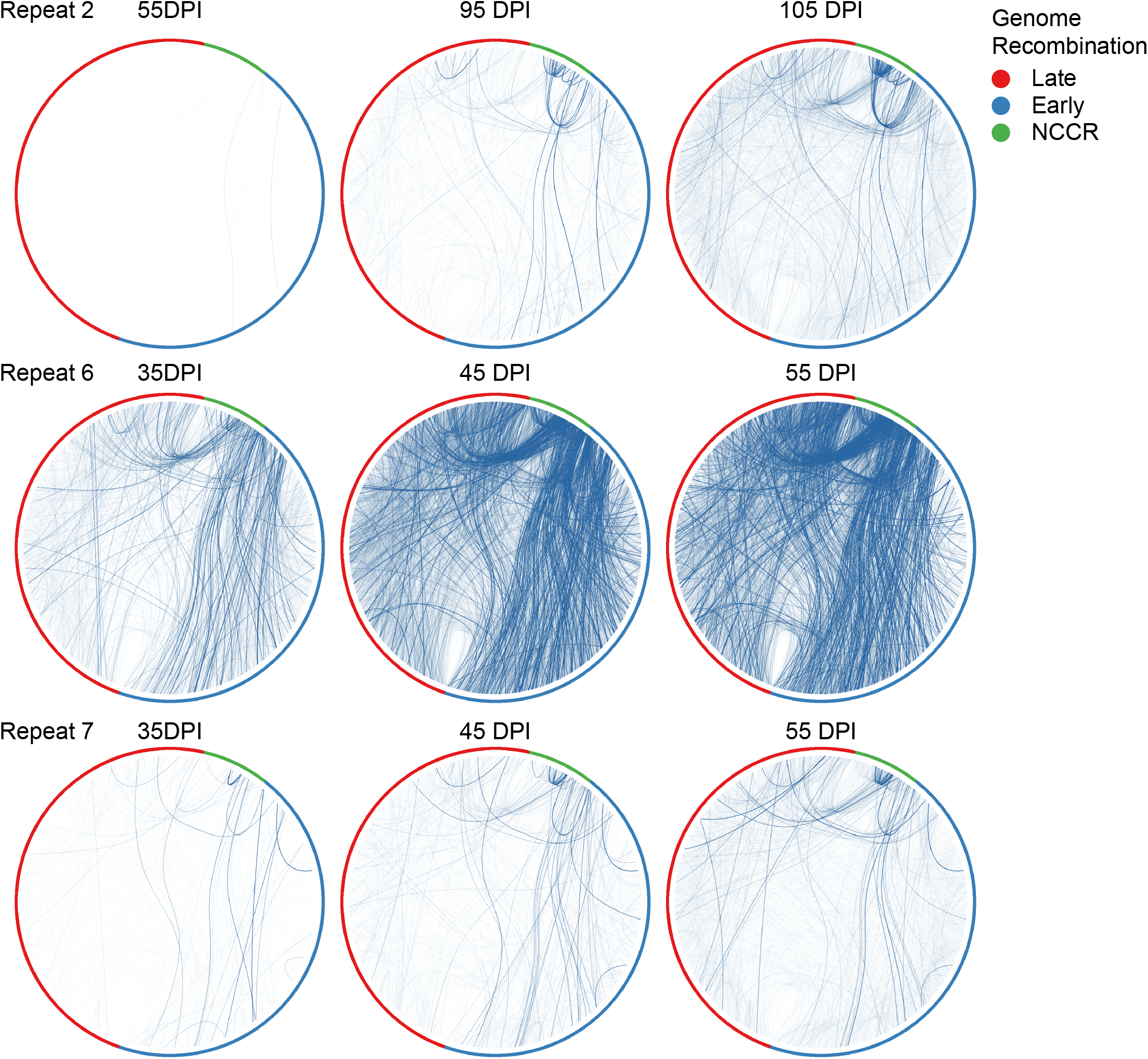
BKPyV genome recombination during persistence and reactivation. Recombination junctions at representative time points from three of the repeats are depicted as lines joining positions on the circular genome as in Figure 3A. The intensity of each line reflects the number of reads corresponding to each junction.

**Fig. S4.**
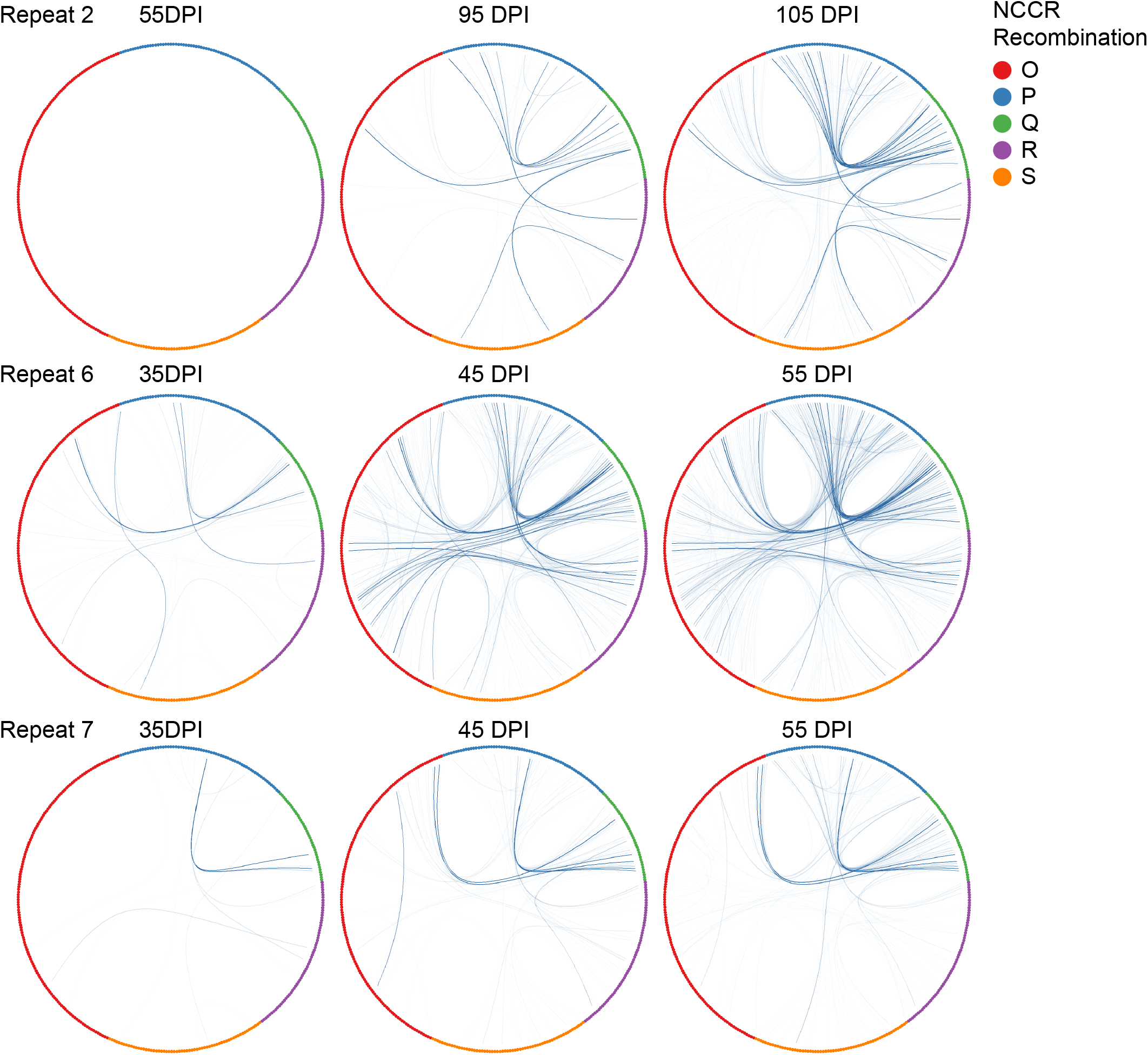
BKPyV recombination within the NCCR during persistence and reactivation. Recombination junctions within the NCCR at representative time points are depicted as in Figure S2 but showing just the NCCR. The intensity of each line reflects the number of reads corresponding to each junction.

**Fig. S5.**
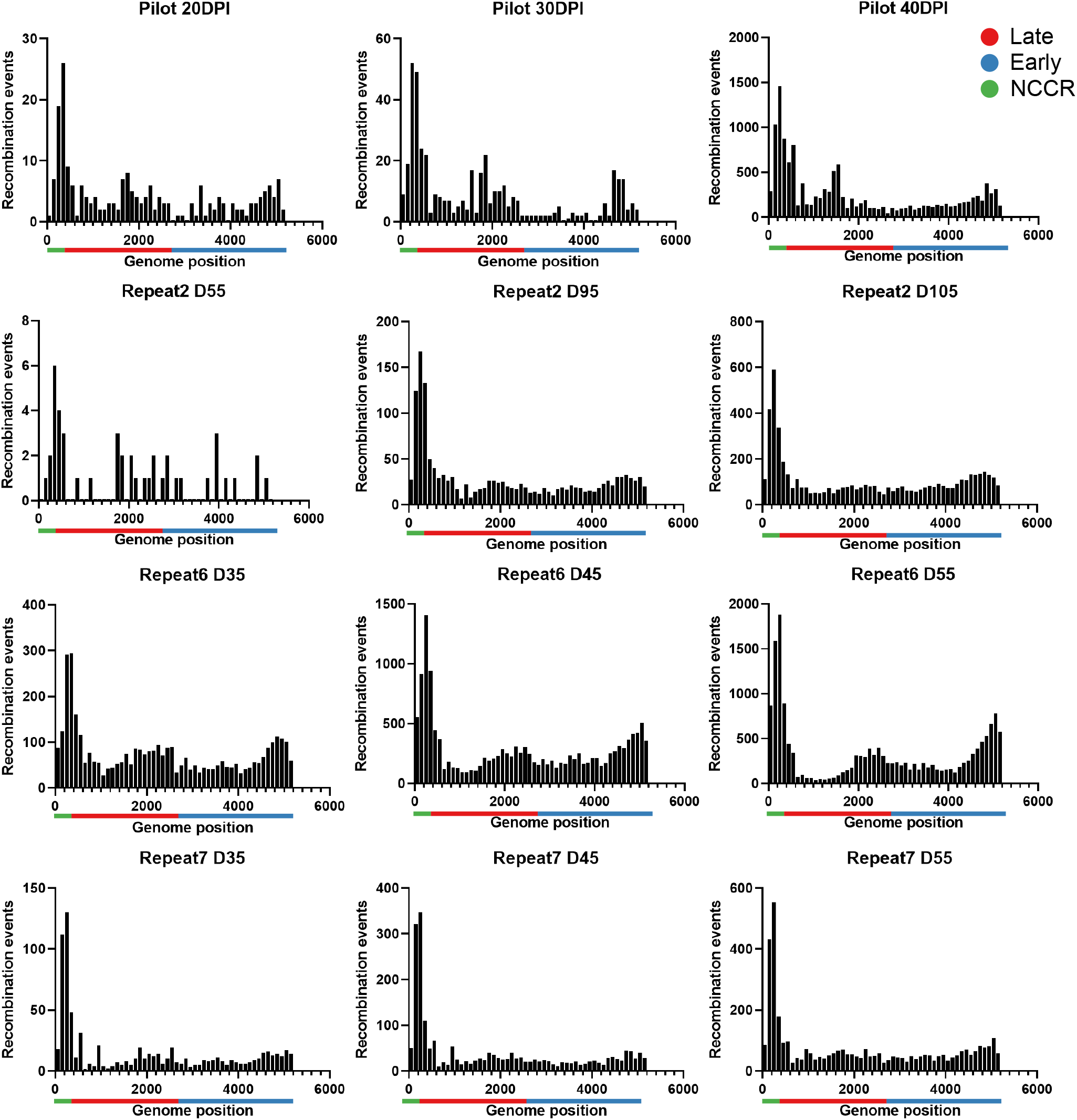
Distribution of unique recombination events. Genome recombination events at representative time points from the pilot and three repeats are illustrated as histograms with each bar representing 100bp. Green bar: NCCR. Red bar: late region. Blue bar: early region.

**Fig. S6.**
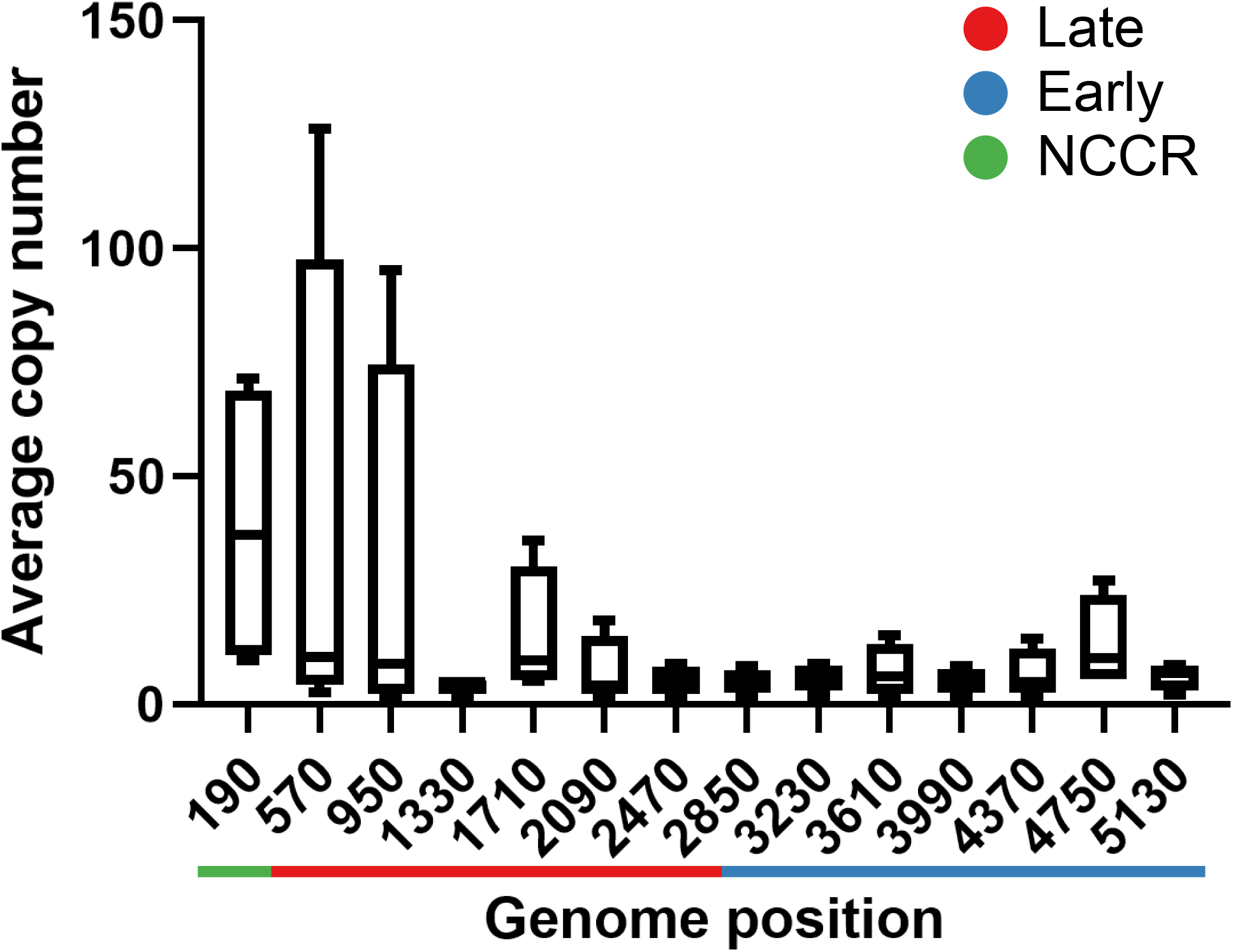
Average copy number of each unique recombination junction. The average copy number was calculated as total recombination read counts/unique recombination counts (deep sequencing reads per unique recombination junction) in each 380 bp segment of the genome. Results from pilot experiment at 40DPI, Repeat 2 105 DPI, Repeat 6 55DPI, Repeat 7 55 DPI are plotted together. Green bar: NCCR. Red bar: late region. Blue bar: early region.

**Fig. S7.**
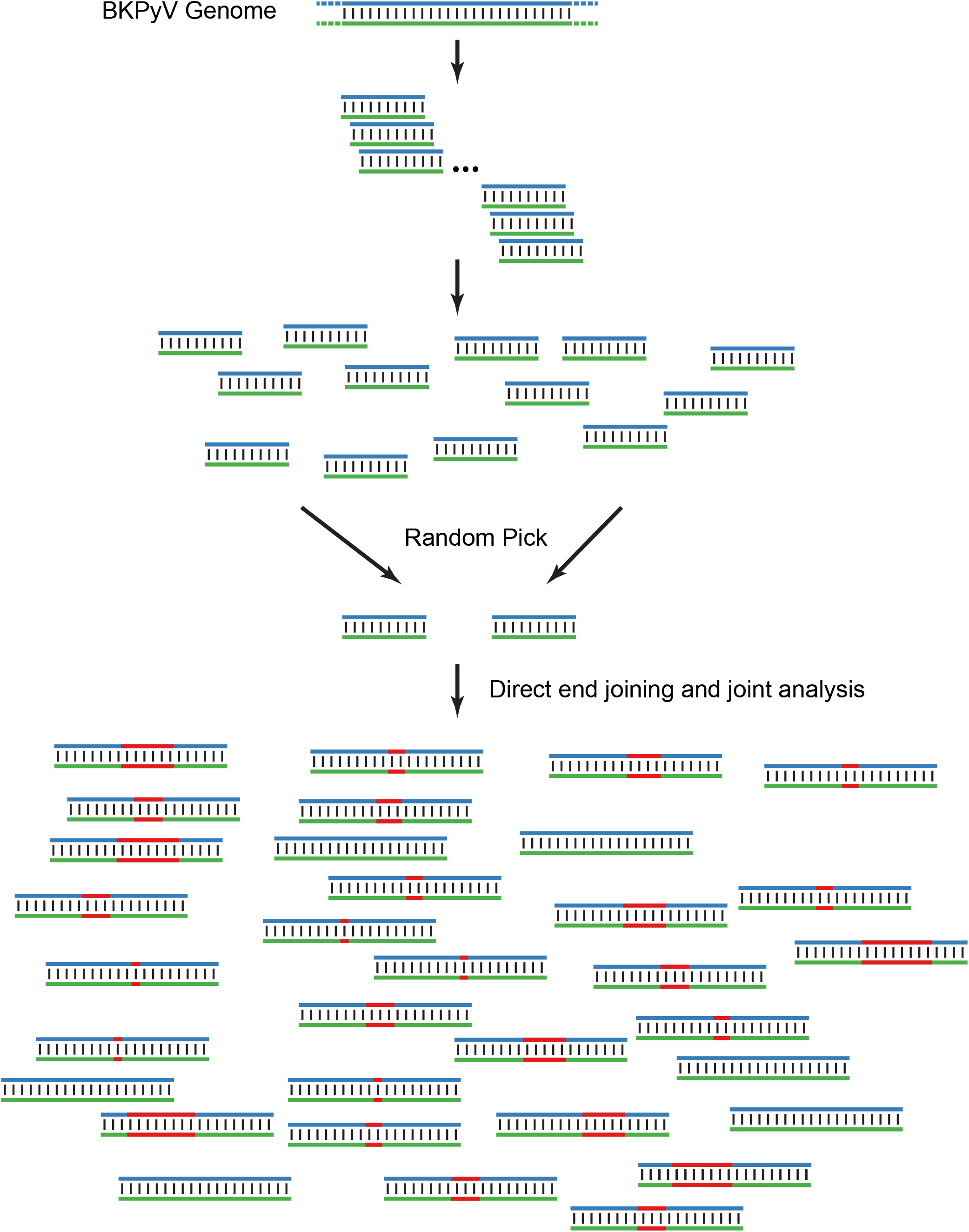
Homology length simulation. See Methods for details.

## References

1. DeCaprio JA, Imperiale MJ, Major EO. 2013. Polyomaviruses. In Knipe DM, Howley P (ed), Fields Virology, 6 ed.

2. Gardner SD, Field AM, Coleman DV, Hulme B. 1971. New human papovavirus (B.K.) isolated from urine after renal transplantation. Lancet 1:1253–7.

3. Padgett BL, Walker DL, ZuRhein GM, Eckroade RJ, Dessel BH. 1971. Cultivation of papova-like virus from human brain with progressive multifocal leucoencephalopathy. Lancet 1:1257–60.

4. Kean JM, Rao S, Wang M, Garcea RL. 2009. Seroepidemiology of human polyomaviruses. PLoS Pathog 5:e1000363.

5. Kardas P, Leboeuf C, Hirsch HH. 2015. Optimizing JC and BK polyomavirus IgG testing for seroepidemiology and patient counseling. J Clin Virol 71:28–33.

6. Jiang M, Abend JR, Johnson SF, Imperiale MJ. 2009. The role of polyomaviruses in human disease. Virology 384:266–73.

7. Broekema NM, Abend JR, Bennett SM, Butel JS, Vanchiere JA, Imperiale MJ. 2010. A system for the analysis of BKV non-coding control regions: application to clinical isolates from an HIV/AIDS patient. Virology 407:368–73.

8. Egli A, Infanti L, Dumoulin A, Buser A, Samaridis J, Stebler C, Gosert R, Hirsch HH. 2009. Prevalence of polyomavirus BK and JC infection and replication in 400 healthy blood donors. J Infect Dis 199:837–46.

9. Olsen GH, Hirsch HH, Rinaldo CH. 2009. Functional analysis of polyomavirus BK non-coding control region quasispecies from kidney transplant recipients. J Med Virol 81:1959–67.

10. Ajuh ET, Wu Z, Kraus E, Weissbach FH, Bethge T, Gosert R, Fischer N, Hirsch HH. 2018. Novel Human Polyomavirus Noncoding Control Regions Differ in Bidirectional Gene Expression according to Host Cell, Large T-Antigen Expression, and Clinically Occurring Rearrangements. J Virol 92.

11. Gosert R, Rinaldo CH, Funk GA, Egli A, Ramos E, Drachenberg CB, Hirsch HH. 2008. Polyomavirus BK with rearranged noncoding control region emerge in vivo in renal transplant patients and increase viral replication and cytopathology. J Exp Med 205:841–52.

12. Bethge T, Hachemi HA, Manzetti J, Gosert R, Schaffner W, Hirsch HH. 2015. Sp1 sites in the noncoding control region of BK polyomavirus are key regulators of bidirectional viral early and late gene expression. J Virol 89:3396–411.

13. Low J, Humes HD, Szczypka M, Imperiale M. 2004. BKV and SV40 infection of human kidney tubular epithelial cells in vitro. Virology 323:182–8.

14. Imperiale MJ, Jiang M. 2016. Polyomavirus Persistence. Annu Rev Virol 3:517–532.

15. Witkin AE, Banerji J, Bullock PA. 2020. A model for the formation of the duplicated enhancers found in polyomavirus regulatory regions. Virology 543:27–33.

16. Zhao L, Imperiale MJ. 2019. Establishing Renal Proximal Tubule Epithelial-Derived Cell Lines Expressing Human Telomerase Reverse Transcriptase for Studying BK Polyomavirus. Microbiol Resour Announc 8.

17. Bennett SM, Jiang M, Imperiale MJ. 2013. Role of cell-type-specific endoplasmic reticulum-associated degradation in polyomavirus trafficking. J Virol 87:8843–52.

18. Zhao L, Marciano AT, Rivet CR, Imperiale MJ. 2016. Caveolin- and clathrin-independent entry of BKPyV into primary human proximal tubule epithelial cells. Virology 492:66–72.

19. Veltri KL, Espiritu M, Singh G. 1990. Distinct genomic copy number in mitochondria of different mammalian organs. J Cell Physiol 143:160–4.

20. Tanaka N, Takahara A, Hagio T, Nishiko R, Kanayama J, Gotoh O, Mori S. 2020. Sequencing artifacts derived from a library preparation method using enzymatic fragmentation. PLoS One 15:e0227427.

21. Bethge T, Ajuh E, Hirsch HH. 2016. Imperfect Symmetry of Sp1 and Core Promoter Sequences Regulates Early and Late Virus Gene Expression of the Bidirectional BK Polyomavirus Noncoding Control Region. J Virol 90:10083–10101.

22. Broekema NM, Imperiale MJ. 2012. Efficient propagation of archetype BK and JC polyomaviruses. Virology 422:235–41.

23. Sowd GA, Li NY, Fanning E. 2013. ATM and ATR activities maintain replication fork integrity during SV40 chromatin replication. PLoS Pathog 9:e1003283.

24. Truong LN, Li Y, Shi LZ, Hwang PY, He J, Wang H, Razavian N, Berns MW, Wu X. 2013. Microhomology-mediated End Joining and Homologous Recombination share the initial end resection step to repair DNA double-strand breaks in mammalian cells. Proc Natl Acad Sci U S A 110:7720–5.

25. Wang H, Xu X. 2017. Microhomology-mediated end joining: new players join the team. Cell Biosci 7:6.

26. Zaragoza C, Li RM, Fahle GA, Fischer SH, Raffeld M, Lewis AM, Jr., Kopp JB. 2005. Squirrel monkeys support replication of BK virus more efficiently than simian virus 40: an animal model for human BK virus infection. J Virol 79:1320–6.

27. Jiang M, Abend JR, Tsai B, Imperiale MJ. 2009. Early events during BK virus entry and disassembly. J Virol 83:1350–8.

28. Jiang M, Zhao L, Gamez M, Imperiale MJ. 2012. Roles of ATM and ATR-mediated DNA damage responses during lytic BK polyomavirus infection. PLoS Pathog 8:e1002898.

29. Zhao L, Imperiale MJ. 2017. Identification of Rab18 as an Essential Host Factor for BK Polyomavirus Infection Using a Whole-Genome RNA Interference Screen. mSphere 2.

30. An P, Saenz Robles MT, Duray AM, Cantalupo PG, Pipas JM. 2019. Human polyomavirus BKV infection of endothelial cells results in interferon pathway induction and persistence. PLoS Pathog 15:e1007505.

31. Assetta B, De Cecco M, O’Hara B, Atwood WJ. 2016. JC Polyomavirus Infection of Primary Human Renal Epithelial Cells Is Controlled by a Type I IFN-Induced Response. mBio 7.

32. Cioni M, Mittelholzer C, Wernli M, Hirsch HH. 2013. Comparing effects of BK virus agnoprotein and herpes simplex-1 ICP47 on MHC-I and MHC-II expression. Clin Dev Immunol 2013:626823.

33. Handala L, Blanchard E, Raynal PI, Roingeard P, Morel V, Descamps V, Castelain S, Francois C, Duverlie G, Brochot E, Helle F. 2020. BK Polyomavirus Hijacks Extracellular Vesicles for En Bloc Transmission. J Virol 94.

34. Verhalen B, Starrett GJ, Harris RS, Jiang M. 2016. Functional Upregulation of the DNA Cytosine Deaminase APOBEC3B by Polyomaviruses. J Virol 90:6379–6386.

35. Justice JL, Needham JM, Thompson SR. 2019. BK Polyomavirus Activates the DNA Damage Response To Prolong S Phase. J Virol 93.

36. An P, Cantalupo PG, Zheng W, Saenz-Robles MT, Duray AM, Weitz D, Pipas JM. 2021. Single-Cell Transcriptomics Reveals a Heterogeneous Cellular Response to BK Virus Infection. J Virol 95.

37. Myhre MR, Olsen GH, Gosert R, Hirsch HH, Rinaldo CH. 2010. Clinical polyomavirus BK variants with agnogene deletion are non-functional but rescued by trans-complementation. Virology 398:12–20.

38. Van Loy T, Thys K, Ryschkewitsch C, Lagatie O, Monaco MC, Major EO, Tritsmans L, Stuyver LJ. 2015. JC virus quasispecies analysis reveals a complex viral population underlying progressive multifocal leukoencephalopathy and supports viral dissemination via the hematogenous route. J Virol 89:1340–7.

39. Addetia A, Phung Q, Bradley BT, Lin MJ, Zhu H, Xie H, Huang ML, Greninger AL. 2021. In Vivo Generation of BK and JC Polyomavirus Defective Viral Genomes in Human Urine Samples Associated with Higher Viral Loads. J Virol 95.

40. Verhalen B, Justice JL, Imperiale MJ, Jiang M. 2015. Viral DNA replication-dependent DNA damage response activation during BK polyomavirus infection. J Virol 89:5032–9.

41. Abend JR, Low JA, Imperiale MJ. 2007. Inhibitory effect of gamma interferon on BK virus gene expression and replication. J Virol 81:272–9.

42. Arad U. 1998. Modified Hirt procedure for rapid purification of extrachromosomal DNA from mammalian cells. Biotechniques 24:760–2.

43. Jiang M, Entezami P, Gamez M, Stamminger T, Imperiale MJ. 2011. Functional reorganization of promyelocytic leukemia nuclear bodies during BK virus infection. mBio 2:e00281–10.

